# Parameter estimation and identifiability analysis for a bivalent analyte model of monoclonal antibody-antigen binding

**DOI:** 10.1101/2022.12.05.519088

**Authors:** Kyle Nguyen, Kan Li, Kevin Flores, Georgia D. Tomaras, S. Moses Dennison, Janice M. McCarthy

## Abstract

1

Discovery research for therapeutic antibodies and vaccine development requires an in-depth understanding of antibody-antigen interactions. Label-free techniques such as Surface Plasmon Resonance (SPR) enable the characterization of biomolecular interactions through kinetics measurements, typically by binding antigens in solution to monoclonal antibodies immobilized on a SPR chip. 1:1 Langmuir binding model is commonly used to fit the kinetics data and derive rate constants. However, in certain contexts it is necessary to immobilize the antigen to the chip and flow the antibodies in solution. One such scenario is the screening of monoclonal antibodies (mAbs) for breadth against a range of antigens, where a bivalent analyte binding model is required to adequately describe the kinetics data unless antigen immobilizaion density is optimized to eliminate avidity effects. A bivalent analyte model is offered in several existing software packages intended for standard throughput SPR instruments, but lacking for high throughput SPR instruments. Existing methods also do not explore multiple local minima and parameter identifiability, issues common in non-linear optimization. Here, we have developed a method for analyzing bivalent analyte binding kinetics directly applicable to high throughput SPR data collected in a non-regenerative fashion, and have included a grid search on initial parameter values and a profile likelihood method to determine parameter identifiability. We fit the data of a broadly neutralizing HIV-1 mAb binding to HIV-1 envelope glycoprotein gp120 to a system of ordinary differential equations modeling bivalent binding. Our identifiability analysis discovered a non-identifiable parameter when data is collected under the standard experimental design for monitoring the association and dissociation phases. We used simulations to determine an improved experimental design, which when executed, resulted in the reliable estimation of all rate constants. These methods will be valuable tools in analyzing the binding of mAbs to an array of antigens to expedite therapeutic antibody discovery research.

**Author summary:** While commercial software programs for the analysis of bivalent analyte binding kinetics are available for low-throughput instruments, they cannot be easily applied to data generated by high-throughput instruments, particularly when the chip surface is not regenerated between titration cycles. Further, existing software does not address common issues in fitting non-linear systems of ordinary differential equations (ODEs) such as optimizations getting trapped in local minima or parameters that are not identifiable. In this work, we introduce a pipeline for analysis of bivalent analyte binding kinetics that 1) allows for the use of high-throughput, non-regenerative experimental designs, 2) optimizes using several sets of initial parameter values to ensure that the algorithm is able to reach the lowest minimum error and 3) applies a profile likelihood method to explore parameter identifiability. In our experimental application of the method, we found that one of the kinetics parameters (*k*_*d*2_) cannot be reliably estimated with the standard length of the dissociation phase. Using simulation and identifiability analysis we determined the optimal length of dissociation so that the parameter can be reliably estimated, saving time and reagents. These methodologies offer robust determination of the kinetics parameters for high-throughput bivalent analyte SPR experiments.

## 3 Introduction

Studying antibody-antigen interactions is crucial for understanding antibody function and potency. Surface Plasmon Resonance (SPR) is a label-free technique enabling characterization of epitope specificities and determination of binding affinities of monoclonal antibodies (mAbs) targeting pathogen surface proteins. Affinity measurements of antibody-antigen binding by SPR are usually carried out by immobilizing the bivalent antibodies (ligand) on the sensor surface and testing the binding of antigens (analyte) in solution. The change in refractive index due to changes in the deposited mass that occurs during analyte binding to- and dissociating from the ligand surface is detected in real time. Typically, when the conditions are optimized, SPR data of antigen binding to immobilized mAbs can be fitted using the 1:1 Langmuir (monovalent) binding model [1, 2, 3, 4].

While immobilizing antibodies allows for rapid generation of affinity data for antigens against multitude of antibodies depending on the throughput of the platform used, there are circumstances where it may be required that antigens be immobilized on the SPR chip. For example, evaluation of binding breadth of antibody candidates to different variants of the antibody targets would require immobilization of different antigens on sensor surfaces to make an array of antigens and test the binding of antibody analytes. If assays are performed in this orientation, the 1:1 Langmuir binding model cannot be used and the bivalency of antibody analytes should be taken into account. Further, even when the antibodies are immobilized, for interactions involving antigens that are dimeric or trimeric, the 1:1 model is not sufficient to describe the kinetics of binding between antibody and antigen. Multivalent binding models, particularly bivalent models when using antibodies as analytes, should be considered in lieu of the 1:1 binding model to describe more complex interactions [5, 6, 7].

Models for bivalent binding kinetics data have previously been developed and studied [8, 9, 10], and commercial software programs exist and have been used to estimate the on- and off-rates of bivalent biomolecular interactions [7, 11, 12, 13, 14]. However these commercial packages have the following limitations: 1) they are designed for low-throughput instruments and for regenerative titration cycles, 2) they do not address issues common in non-linear optimization such as local minima and parameter identifiablity. Both limitations are elaborated below.

In a typical SPR binding kinetics assay, the ligand is first immobilized onto the SPR chip either through permanent immobilization, such as amine-coupling or streptavidin capturing, or through non-permanent capture using surface molecules that show strong affinity to the ligand. Next, a titration is performed to detect the association and dissociation of analyte to immobilized ligand at multiple concentrations, typically sequentially from low to high. Analysis of kinetics traces at multiple concentrations enhance the accuracy of kinetics parameter estimation. During each titration cycle, one first flows buffer over the chip to obtain baseline, followed by an association phase where the analyte is flowed over the chip and a dissociation phase where only buffer is flowed.

Before the start of each cycle, the chip surface may be regenerated typically using a buffer with extreme pH values or high concentration of salt. If the ligand is permanently immobilized and can withstand extreme conditions, regeneration can rapidly dissociate the analyte from ligand and frees the ligand on the chip. If the ligand is non-permanently captured, regeneration can dissociate the analyte-ligand pairs from the ligand-capturing molecules, enabling re-capturing of the ligand before the next titration cycle. However, regeneration is not always feasible, permanently immobilized ligands are often sensitive to extreme conditions; the re-capturing of ligand in every cycle could also lead to longer experiment time and higher reagent consumption. In these cases, no regeneration will be implemented, and therefore the SPR chip is not completely free of bound analyte when the next cycle starts.

Whether we are studying 1:1 binding, bivalent or multivalent interactions, we obtain the kinetics parameters by fitting the SPR data to a system of ordinary differential equations (ODEs), using an optimization algorithm that compares the model’s predictions to the data and finds the ‘best’ parameters by minimizing the error in predicted outcome versus observed.

The so-called ‘local minima’ problem is common in non-linear optimization. One can think of the problem as a very ‘bumpy’ curve. Starting at an initial value supplied by the user, the algorithm works in small steps to find the lowest bump. Depending on where one starts, it is possible to get “stuck” in a part of the curve that is not the lowest part. As an example, Fig 1 illustrates a hypothetical case where the error (or sum of squares error) is a function of a single parameter value. For a robust fit, we perform parameter estimation at multiple sets of initial guesses, i.e. optimize using several sets of initial parameter values that cover wide ranges of kinetics parameters, and record the optimized parameters with the lowest error. This approach ensures that recorded optimized parameters give the lowest possible sum of squared for error (purple triangle in Fig 1).

**Fig 1:**
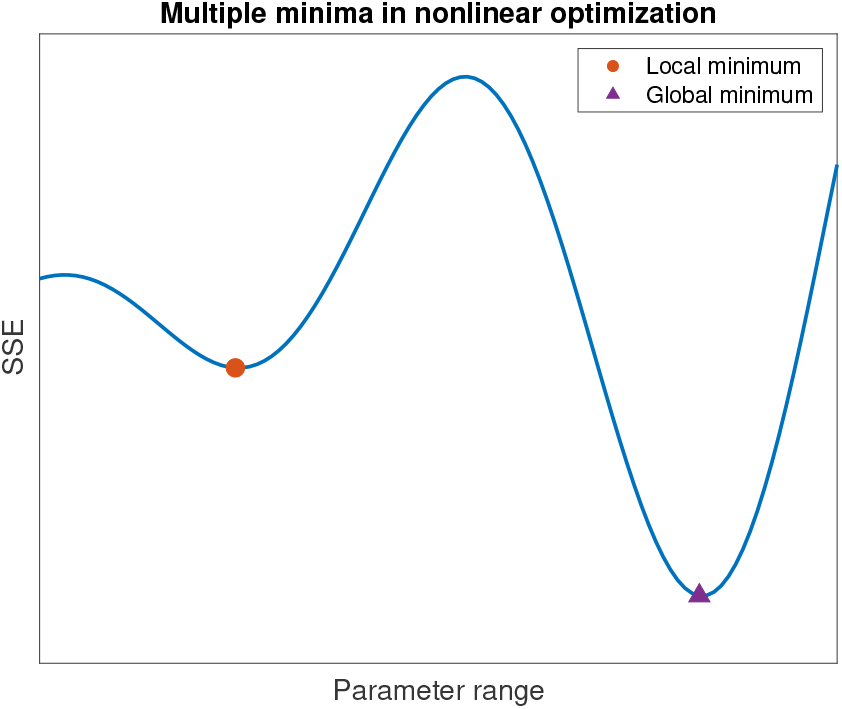
Illustrative example of a multiple minima problem. Illustrative example of a multiple minima problem that is common in nonlinear optimization is shown. The blue curve represents the sum of squares error (SSE) as a function of different values of a parameter. The orange dot is a local minimum and the purple triangle is the global minimum.

Unlike the local minima problem, non-identifiability happens when one or more parameters cannot be reliably estimated because different values of the parameter(s) lead to the same (or numerically similar) values in the error function. Identifiability issues can arise when a model has many parameters compared to observed data or when there are unobserved states. Mathematical methods for parameter identifiability analysis have been developed, described, and improved upon in the literature [15, 16] and have been applied in many biomathematical models, for example in epidemiology [17, 18, 19] and oncology [20, 21].

We explain these problems of local minima and non-identifiability in detail in the body of the paper. For the problem of parameter non-identifiability, we go further than the usual mathematical analysis and use simulation to guide experimental design. This computational step is especially important for improving resource and time efficiency during the data collection step while ensuring that there is sufficient information in the data to fit the model.

## 4 Materials and methods

We first provide an overview of our experimental procedure in the form of a schematic. As illustrated in Fig 2, we start with model refinement and code development for the bivalent analyte binding model, then experimental data collection. This is followed by the parameter estimation step and parameter identifiability analysis using the log profile likelihood [15]. If a parameter is found to be practically non-identifiable, before collecting new experimental data, we simulate synthetic data and find a new data collection scheme that will give us more information about the parameter. Applying parameter identifiability analysis to synthetic data gives us an estimate of the length of the relevant phase necessary to identify all parameters. In the case we are presenting, we extended the dissociation phase to optimize the identifiability of one of the dissociation constants. Then we collect new experimental data using the optimized data collection scheme. If parameter estimation on the new experimental data confirms that all parameters are identifiable, we proceed to report the estimates.

**Fig 2:**
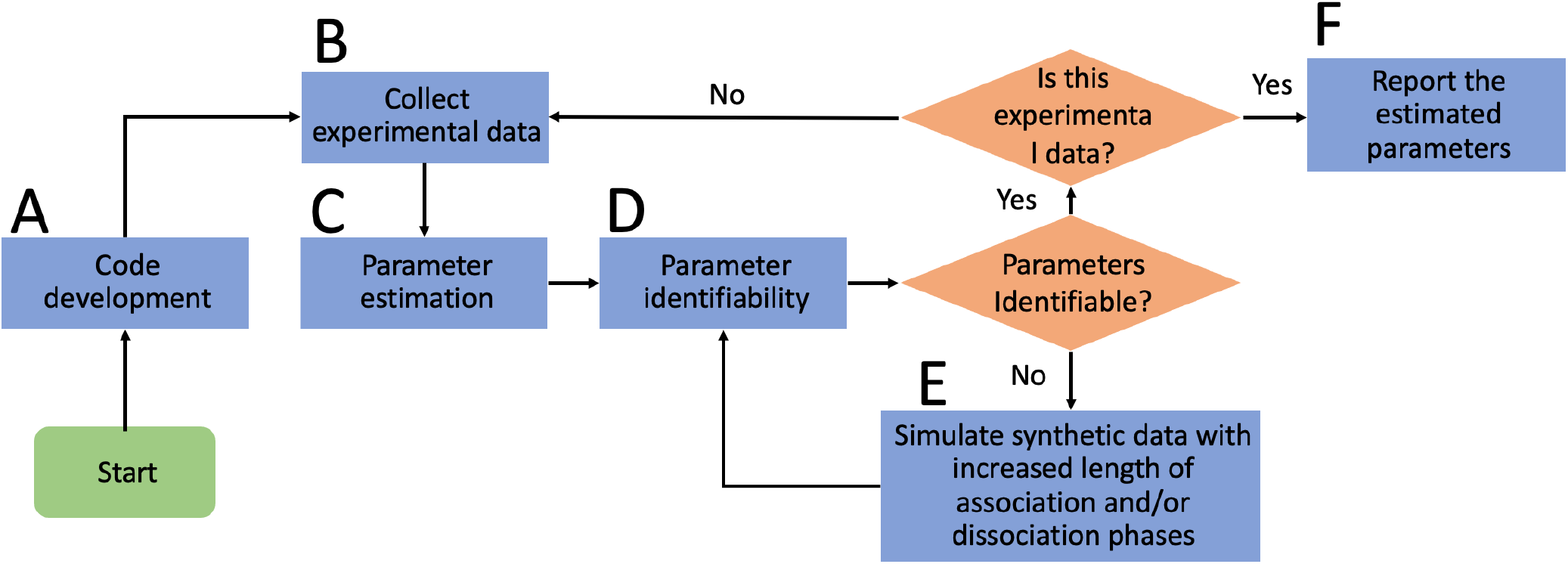
Schematic for the experimental procedure. Schematic depicting the step-by-step process for reliable determination of the kinetics parameters for the bivalent analyte model. (A) First, we start with the code development and strategy for parameter estimation for data collected in high throughput fashion. (B) Next, we collect experimental data. (C) We then obtain estimates of the parameters using non-linear optimization. (D) After obtaining the estimates, we examine the identifiability of the parameters. (E) If a parameter is found to be non-identifiable, we perform simulations to find the optimal data collection scheme in which that parameter is identifiable. After finding the optimal scheme, we restart the process from step (B) and confirm that the parameters are identifiable. (F) Finally, we report the estimated parameters.

### 4.1 Experimental data collection

The binding kinetics measurements of HIV antibody constructs were done on the Carterra LSA platform using HC30M sensor chips (Carterra) at 25°C. Two microfluidic modules, a 96-channel print-head (96PH) and a single flow cell (SFC), were used to deliver liquids onto the sensor chip.

The chip was first activated by 100 mM N-Hydroxysuccinimide (NHS) and 400 mM 1-Ethyl-3-(3-dimethylaminopropyl) carbodiimide hydrochloride (EDC) (Cytiva, mixed 1:1:1 with 0.1 M MES buffer at pH 5.5) for 600 seconds, followed by direct immobilization of CH505 transmitted founder (T/F) gp120 [22] (in 10 mM Sodium Acetate at pH 4.5) at multiple concentrations for 600 seconds using the 96PH. Unreactive esters were then quenched with a 600 seconds injection of 1 M ethanolamine-HCl at pH 8.5. The running buffer was 10 mM MES buffer at pH 5.5 with 0.01% Tween-20, and each concentration of CH505 T/F gp120 was immobilized onto 24 separate spots of the same chip. Unless specified above, the steps were done using the SFC. CH505 T/F gp120 was produced by Duke Human Vaccine Institute Protein Production Facility as described earlier [22] and further purified by size exclusion chromatography for monomeric gp120.

A two-fold dilution series of the CH31 monoclonal antibody, an HIV-1 mAb with CD4-binding site specificity [23], was prepared in 1× HBSTE (10 mM HEPES pH 7.4, 150 mM NaCl, 3 mM EDTA and 0.01% Tween-20) buffer. The highest concentration was 150 μg/ml (1.0 μM). CH31 at different concentrations was then injected using SFC onto the chip surface from the lowest to the highest concentration without regeneration, including 8 injections of buffer before the lowest non-zero concentration for signal stabilization. For each concentration, the data collection time-length for baseline and association was 120 seconds and 300 seconds, respectively; the standard time-length for dissociation was 600 seconds and the extended time-length for dissociation was 1800 seconds. For all assays the running buffer for titration was 1X HBSTE.

The titration data collected were first pre-processed in the Kinetics (Carterra) software, including reference subtraction using spots with no immobilized biomolecules, buffer subtraction using the last zero-concentration cycle and data smoothing. The data were then imported into Excel from the Kinetics package. 14 spots that show sensorgrams with good dose dependence and least amount of noise were down-selected for bivalent model analysis.

In Fig 3b, we show a representative sensorgram of the bivalent analyte interaction between CH31 and CH505 gp120. Each sensorgram shows the data from a non-regenerative titration, constituted of 10 binding response curves where CH31 concentration increases two fold from each previous cycle. The vertical solid and dashed lines separate the response curves from left to right into: baseline, association, and dissociation. Before parameter estimation, we performed baseline correction using the baseline data and then chose the 5 ‘best’ concentrations (i.e., 6.25 × 10^-8^, 1.25 × 10^-7^, 2.5 × 10^-7^, 5 × 10^-7^, and 10^-6^ M) for fitting kinetic constants. The 5 concentrations chosen are in the linear range of the dose response (log_10_ concentration versus end of association response) graph as shown in Fig 3c.

**Fig 3:**
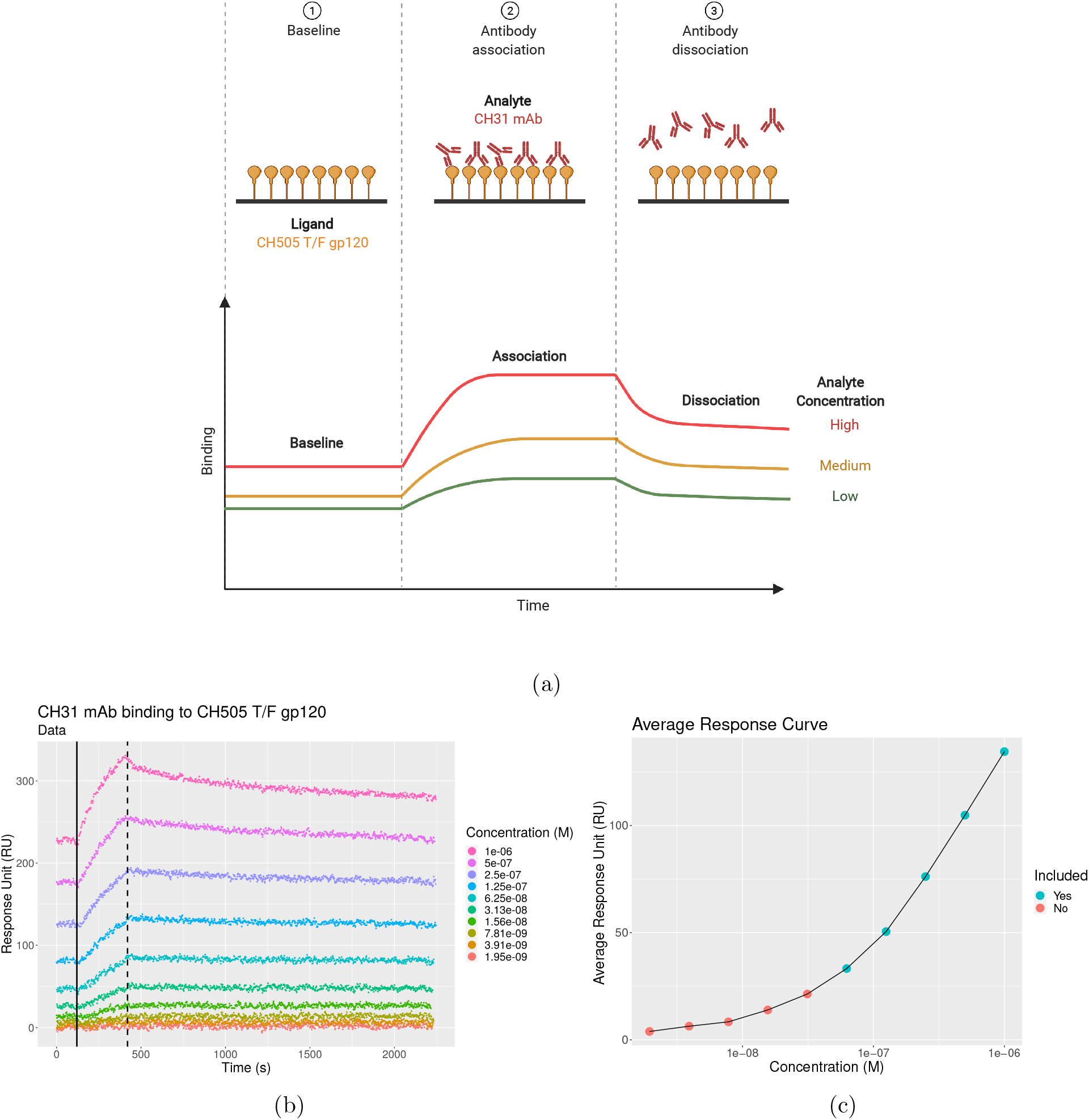
Bivalent analyte association and dissociation processes with a representative sensorgram. **(a)** Bivalent analyte association and dissociation processes. (1) During the baseline phase, the response on the sensor is stabilized. (2) During the association phase, analyte solution that contains CH31 mAb is flowed over. The analytes start associating to the ligands (CH505 T/F gp120) on the sensor. This results in an increase in response. During the dissociation phase, buffer solution with no analyte is flowed allowing the analytes to begin dissociating from the ligands, resulting in decreasing response. **(b)** Stacked sensorgram of a bivalent analyte CH31 mAb binding to a transmitted/founder CH505 HIV-1 envelope glycoprotein gp120 antigen (ligand) are shown. The 10 **dotted** curves correspond to 10 different concentrations of CH31 in the solution with the concentration values being shown in the legend. The left vertical **black solid** line separates the baseline and association, while the right vertical **black dashed** line separates the association and dissociation steps. **(c)**A plot of binding response averaged at the end of association step is shown as a function concentration of CH31 mAb. The data corresponding to the **Yes** concentrations are chosen for fitting kinetic constants. The data for **No** concentrations are excluded.

### 4.2 Bivalent analyte binding model

The simplest model for binding kinetics is the 1:1 Langmuir model that can be described by the following reaction:

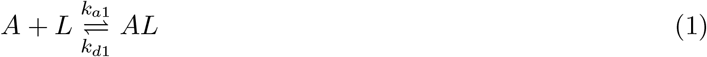

where *A* is the analyte in the solution, and *L* is the immobilized ligand on the sensor. This model assumes that one analyte only binds with one ligand to form the complex *AL* with association rate constant *k*_*a*1_.

The complex *AL* can dissociate into *A* and *L* with dissociation rate constant *k*_*d*1_.

In case of antibody-antigen interaction, Whether the bivalent antibody is immobilized as ligand or used as analyte determines the appropriate binding model (1:1 or bivalent analyte model). SPR measures the change in mass at the chip surface, such that the response is proportional to the concentration of analyte bound to the surface. Therefore when the antibody is immobilized, response increases for each binding site bound to analyte, so that there is a 1:1 interaction between each bound analyte and binding *site* on the ligand. In contrast, when bivalent antibody is used as the *analyte*, the binding of the second antibody arm to an immobilized ligand does not result in further change of response, so that there can be a decrease of free ligand molecules with no change in response. This violates the 1:1 assumption and necessitates a more complicated mathematical model. Fig 4 illustrates how the binding modes differ depending on the whether the antibody or antigen is immobilized.

**Fig 4:**
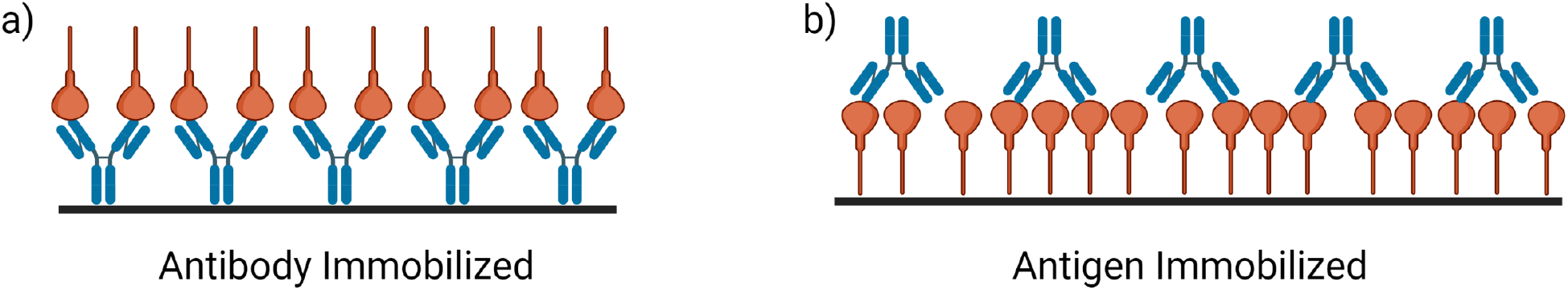
The orientation of binding impacts kinetics characteristics. Cartoon illustration of binding modes when either antibody or antigen is immobilized. The scenario of antibody being immobilized is shown in a), where 1:1 binding interaction can be assumed. The scenario of antigen being immobilized is shown in b), where bivalent model is needed.

When studying antibody-antigen binding, most assay designs allow the antibody to be immobilized and the antigen to be the analyte. However, certain assay designs require the antibody to be the analyte, and analysis by bivalent model is needed. Bivalent analyte binding kinetics can be represented using the following two reversible reactions:

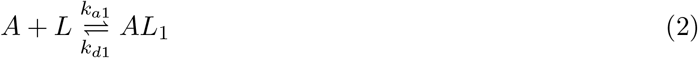

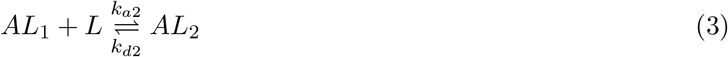

where *A* is the bivalent analyte in the solution, and *L* is the immobilized ligand on the sensor. In this study, *A* and *L* represent CH31 mAb and gp120 of HIV-1, respectively. The complex *AL*_1_ is formed when a ligand binds with one arm of an analyte at the association rate *k*_*a*1_. *AL*_1_ can revert back to *A* and *L* at the rate *k*_*d*1_. In the second reaction, the remaining arm of AL_1_ can associate to- and dissociate from another ligand at the association and dissociation rates, *k*_*a*2_ and *k*_*d*2_, respectively. These processes are further illustrated in Fig 3a. It is important to note that we are unable to observe the difference between [*AL*_1_] and [*AL*_2_] - a detail that can lead to parameter non-identifiability.

During the data collection process, at the beginning of association phase, the solution containing analyte flows through the sensor for association. In the dissociation phase, the buffer solution without analyte flows through the sensor allowing the bound analyte to dissociate. Therefore, we separate the model into two sub-models: association phase and dissociation phase. The details of the association model is described in Section 4.2.1. We describe the dissociation phase model in Sections 4.2.2.

#### 4.2.1 Association phase model

For the association phase, one can derive a bivalent analyte model that consists of ordinary differential equations (ODEs) from the reactions (2) and (3). In this study, we employ a model that is used by commercial software, ForteBio [24] and Biacore [25]. The model is described by the following equations:

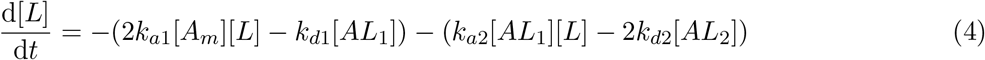

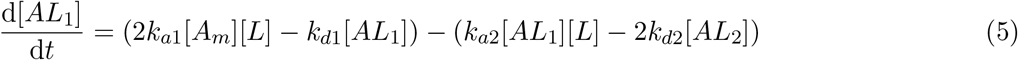

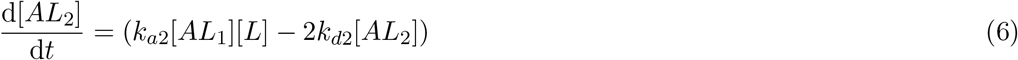

where the [*A_m_*] represents the analyte concentration in the solution and is assumed constant as analyte is continually supplied. [*L*], [*AL*_1_], and [*AL*_2_] represent the concentrations of free ligand, analyte-ligand complex, and analyte-two-ligand complex, respectively. The first two terms of of each Eqs. (4) and (5) are derived from the Eq. (2). The concentration of free ligand [*L*] decreases due to the formation of [*AL*_1_] when a free ligand binds with one of an analyte. The factor of two in the first term of Eq. (4) accounts for the fact that the free ligand can bind at either arm of the analyte. On the other hand, the concentrations of free ligand increases proportionally to the decomposition of *AL*_1_. In the first reaction (Eq. 2), the change in concentration of analyte-ligand, [*AL*_1_], is opposite to the change of [*L*]. As the concentration of free ligand increases or decreases, the concentration of analyte-ligand complex decreases or increases.

This model assumes a two-step process, where the binding and unbinding have to be occur in order. This means *AL*_2_ cannot be formed before the formation of *AL*_1_. Similarly, *AL*_1_ cannot be dissociated before the dissociation of AL2. These assumptions explain the remaining terms in Eqs. (4)–(6). The concentration of analyte-two-ligand complex ([*AL*_2_]) increases proportionally to both concentrations of the free ligand and the analyte-ligand. The analyte-two-ligand complex decompose into an analyte-ligand complex and a ligand. Note that there are again multiple factors of two in these terms to account for the fact that the bound analyte can unbind with either of its arms.

In this high throughput experiment, the SPR chip surface is not regenerated between concentrations of analyte. If the surface were regenerated, we could set [*AL*_1_]_0_ = [*AL*_2_]_0_ = 0. Without regeneration, the concentrations [*AL*_1_]_0_ and [*AL*_2_]_0_ are unknown. To address this, we extrapolate back in time by estimating an initial time adjustment, *t*\*, and compute the initial time *t*_0_ for each concentration as follows:

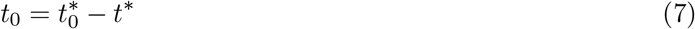

where 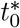 is the initial starting time of the experiment. We then can assume that at this adjusted time *t*_0_, [*AL*_1_]_0_ = [*AL*_2_]_0_ = 0.

The initial amount of free ligand [*L*]_0_ is also unknown. We fit this as a parameter for each concentration.

#### 4.2.2 Dissociation phase model

During the dissociation phase, the buffer containing no analyte is flowing through the cell, allowing the analyte to dissociate from the ligand. In this model, we assume that the association rate constants, *k*_*a*1_ and *k*_*a*2_, are negligible and set them to zero. This means there is no rebinding of the analyte to the ligand. Therefore, the reaction equations for the dissociation phase take the form of two decomposition reactions:

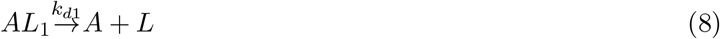

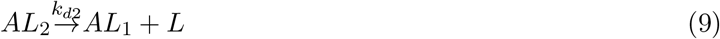

Corazza *et al*. [7] also assumed strict decay of the bound species, but used a sum of exponential model with two parameters, *k*_*d*1_ and *k*_*d*2_. Here, rather than using a phenomenological model, we derived a mechanistic model that consists of the following ODEs:

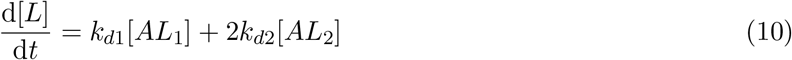

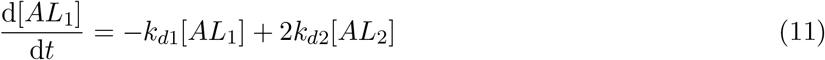

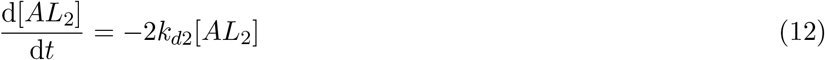

As previously stated in Section 4.2.1, the model follows the assumptions of a two-step process. However, unlike the association phase, during the dissociation phase, we assume the concentration of the analyte-two-ligand complex only decreases over time as *AL*_2_ decomposes into *AL*_1_ and *L*. Each analyte-single-ligand complex, *AL*_1_, is then decomposed into an analyte, *A*, and a ligand, *L*. By using a system of ODEs for the dissociation phase, we make the assumption that *AL*_2_ cannot directly decompose into an *A* and two *L* without decomposing into an *AL*_1_ and an *L* first. Further, we assume that *AL*_1_ cannot rebind to form *AL*_2_. Combining Eqs. (4)–(6) for association model with Eqs. (10)–(12), we have the bivalent analyte model.

### 4.3 Parameter estimation

Unlike the 1:1 Langmuir model, the bivalent analyte model is comprised of nonlinear ODEs. While it is possible to find the analytical solution for the 1:1 Langmuir models, it is not possible to find the analytical solutions for the non-linear bivalent analyte model. Instead, we solve for the approximate solutions by integrating the ODEs numerically. In this work, we use the function ode from an R package called `deSolve` [26], to numerically approximate the solution of the model. Additionally, we use the nonlinear least squares function, `nls.lm`, from the package `minpack` [27], to fit the data using the Levenberg-Marquardt (LM) algorithm [28]. This algorithm is also used in the commercial software Biacore [25]. The `LM` algorithm estimates the parameters by minimizing the sum of squared error (SSE):

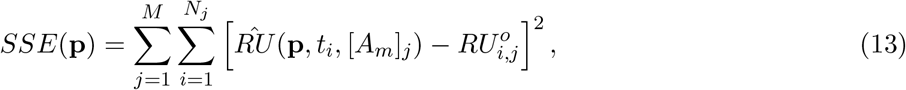

where 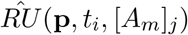 and 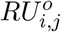, respectively, are model output response unit (RU) and observed response unit in the data at the *i*th time for the *j*th analyte concentration. *N_j_* is the total number of data points observed for the jth concentration and *M* is the total number of concentrations being used during fitting. The vector **p** represents a vector of parameters to be estimated. As explained in Section 4.1, for each data set, we fit the kinetics data for the five concentrations of CH31 at which responses vary in a concentration dependent manner . To improve confidence in parameter estimation of the rate constants (e.g., *k*_*a*1_, *k*_*d*1_, *k*_*a*2_, and *k*_*d*2_), we fit all five concentrations simultaneously, i.e., *M* = 5. This is referred to as a ‘global fit’ and has been shown to yield more robust and reliable results [24]. In addition to the global parameters, each analyte concentration also has a set of local parameters, *R*max_*j*_ and 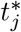, with *j* = 1,…, 5. *R*max’s are the responses associated with maximum analyte bound to the surface, which is proportional to the maximum free ligand concentration on the sensor, [*L*]_0_. Because we are using non-regenerative titration data, we also need to fit an initial time adjustment, 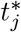, for each concentration.

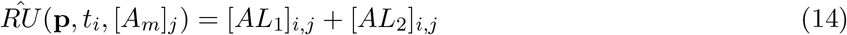

We estimated 14 parameters for each data set: 4 global parameters (*k*_*a*1_, *k*_*d*1_, *k*_*a*2_, and *k*_*d*2_) and 10 local parameters ([*L*]_01,…05_ and 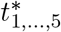). The summary details of all parameters for each model are shown in Table 1. We used a constrained optimization where the lower bound of all parameters was zero.

**Table 1:** Summary of parameters for the bivalent analyte model.

| Parameter | Description | Unit | Fit |
| --- | --- | --- | --- |
| $k_{a1}$ | First binding rate constant | $M^{-1}s^{-1}$ | Global |
| $k_{d1}$ | First unbinding rate constant | $s^{-1}$ | Global |
| $k_{a2}$ | Second binding rate constant | $RU^{-1}s^{-1}$ | Global |
| $k_{d2}$ | Second unbinding rate constant | $RU^{-1}s^{-1}$ | Global |
| $R_{\max}$ | Response associated with the maximum analyte bound to ligand; proportional to $[L]_0$ | $RU$ | Local |
| $t^*$ | Initial time adjustment for non-regenerative titration | $s$ | Local |

### 4.4 Parameter identifiability

Parameters are not identifiable when the global minimum error occurs at multiple values of one or more parameters, i.e., changing the parameter(s) does not effect the error. Parameter identifiability analysis allows modelers to examine whether parameters can be uniquely estimated given the model and the data. There are many reasons for which parameters cannot be uniquely estimated. Non-identifiable parameters may arise when the model is highly complicated, for example, when the number of parameters is too large or when there are too many unobservable states. When the model is the source of non-identifiability, it is called structurally non-identifiable. In this case, some of the model parameters are functions of the others, and as such may vary freely without changing the model output. Fortunately, our model is simple enough that if such structural problems were present, they would be apparent - the model parameters are clearly independent of one another. Therefore, we will not discuss this type of identifiability issue further, but refer the reader to see [17, 18, 20, 19, 21, 29] for more details.

Another source of non-identifiability can be due to the presence of noise or not having enough information in the data. This is called practical non-identifiability. When dealing with practical non-identifiability, one potential approach is to reduce the noise level during the data collection process. However, such an approach is often not possible in practical settings. Rather than trying to reduce the noise level, a more realistic approach is to collect additional data so there is sufficient information for the algorithm to uniquely estimate parameters [18, 20, 29].

#### 4.4.1 Identifiability analysis using the profile likelihood

One of the methods for identifiability analysis is the profile likelihood-based confidence intervals [15]. The profile likelihood creates a profile for each parameter across a reasonable range of values. Given the data, the negative log likelihood function is defined as:

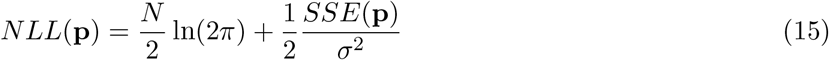

where *NLL*(**p**) is the negative likelihood while **p** is the vector of model parameters. *N* is the number of measurements in the data. In addition, *SSE*(**p**) is computed using Eq. 13. *σ* is the measurement error. We assume that *σ* is known, is the same for all measurements, and can be approximated using residual errors. Furthermore, the minimum of the likelihood is independent of the constant number, 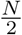 ln(2*π*). Therefore, minimizing the negative likelihood function *NLL* is equivalent to minimizing the function *SSE*. For each parameter *p_j_*, the profile likelihood *PLL_j_* is computed using the following function [21]:

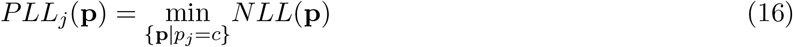

where *c* is a fixed value for *p_j_* within a predefined reasonable range. Simply put, to compute a profile likelihood for parameter *p_j_*, we fix *p_j_* across a range of values with [min(*p_j_*), max(*p_j_*)]. Then, we perform parameter estimation as described in Section 4.3 for all parameters except *p_j_* for each fixed value of *p_j_*. Finally, we compute *NLL* using the estimated parameters (with fixed *p_j_*). The computed *NLL*’s across all fixed values of *p_j_* form a profile likelihood for *p_j_*. The pseudocode to compute profile likelihood is described in Algorithm 1 (in S1 Appendix).

In Fig 5a, we illustrate an example of a practically non-identifiable parameter, where the profile is flat on one side. On the other hand, when the profile likelihood is nonflat on both sides, as shown in Fig 5b, the parameter is practically identifiable. To determine the flatness of the profile, we need to compute the flatness threshold. The threshold is computed as follows [21]:

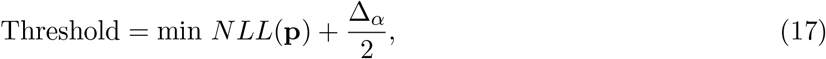

where Δ_*α*_ = *χ*^2^(*α, df*), is the *α*-quantile for the *χ*^2^-distribution. Since we are computing the upper 95% confidence threshold, we choose *α* = 0.95. According to Raue *et al.*, the degree of freedom, *df*, should be either 1 or #p, with #p being the number of parameters to be estimated [15]. To compute the threshold for each individual parameter, the degrees of freedom must be *df* = 1 [15]. We compute a joint threshold for all parameters, so we choose *df* = #p [15].

**Fig 5:**
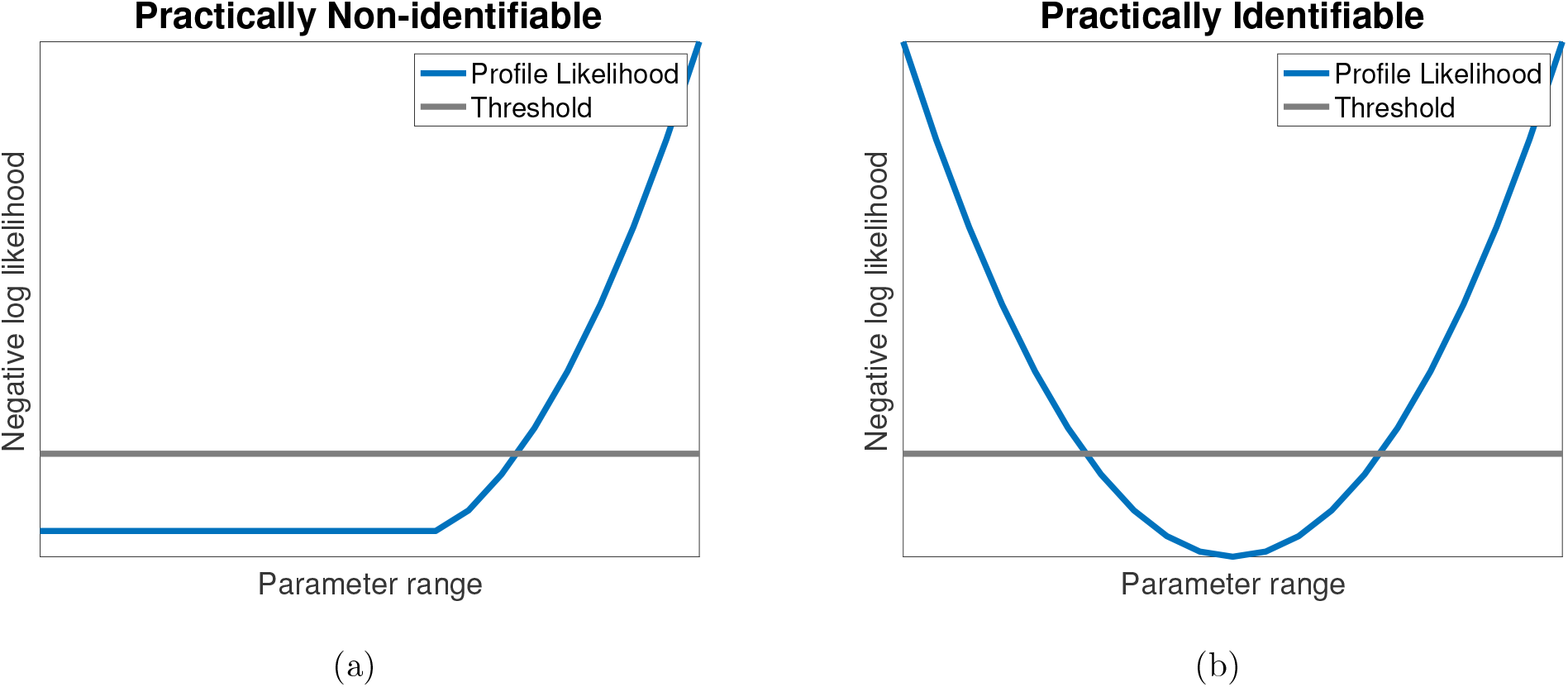
Illustrative examples practically non-identifiable and practically identifiable. Illustrative example profile likelihood for: **(a)** a practically non-identifiable parameter, and **(b)** a practically identifiable parameter. In sub-figure (a), comparing to the threshold (**gray** line), the profile likelihood (**blue** curve) for a structurally non-identifiable parameter are completely flat on both sides. For a practically non-identifiable parameter, as shown sub-figure (b), the profile likelihood is flat on one side and manifests an *L*-shape curve. On the other hand, in sub-figure (c), a practically identifiable parameter has a bowl-shape profile likelihood with a clear minimum.

#### 4.4.2 Simulated data for identifiability analysis

If a parameter is practically non-identifiable for a given data set, additional data is often required to improve parameter identifiability. It is possible to collect experimental data multiple times to incrementally include more temporal points each iteration, but this is certainly not optimal in terms of time and resource efficiency. In previous studies, profile likelihood method has been applied to parameter identifiability analysis to simulated synthetic data of a vector-borne disease model [18, 19] and to a pharmacodynamics model [30]. This approach is especially advantageous because we only need to collect experimental data at most twice.

## 5 Results

### 5.1 Kinetics parameters are not reliably estimated with standard length of dissociation phase

In this section, we present the results of fitting the bivalent analyte model using the standard length of dissociation phase, i.e., 600 seconds. For each data set, we performed global fitting to obtain a more robust result using five different concentrations (6.25 × 10^-8^, 1.25 × 10^-7^, 2.5 × 10^-7^, 5 × 10^-7^, and 1 × 10^-6^ M) of CH31 mAb. In addition, to avoid reporting estimated parameters at local minimum, we performed parameter estimation at 81 different initial guesses for each data set. In Fig 6, we show the fitting result for one example data set (see S1 Fig for all 14 replicates). Each colored scatter curve represents the data for each concentration, while the five solid black curves are the model fitting results. The association and dissociation phases are separated by the black vertical dashed line. Although there are some fitting discrepancies at the end of the association phase for the highest concentration (1 × 10^-6^ M), the bivalent binding model fits the data well (Fig 6).

**Fig 6:**
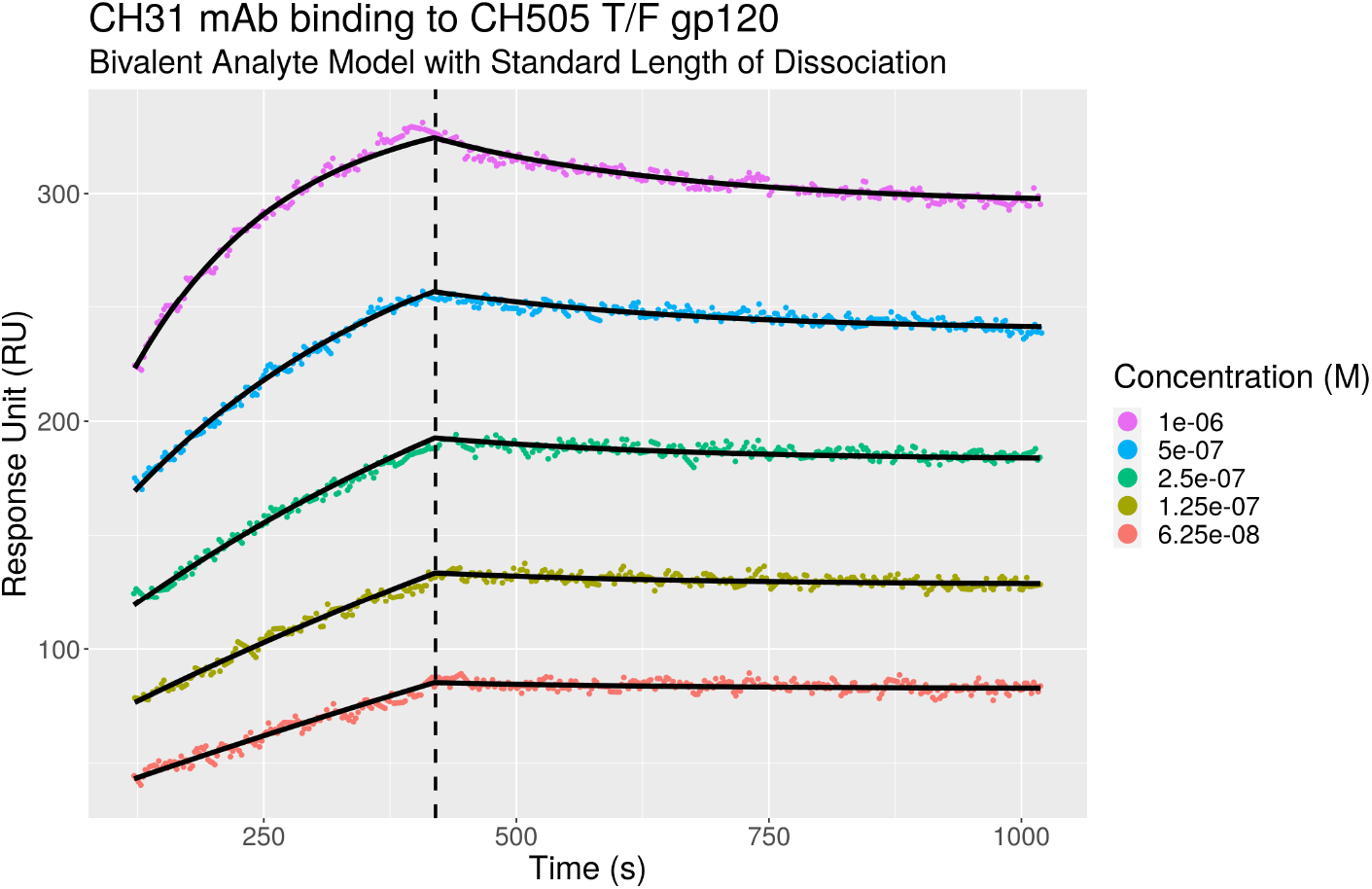
A representative bivalent analyte model fitting result with standard length of dissociation. Bivalent analyte model fitted sensorgrams of CH31 mAb binding to CH505 T/F gp120 are shown. The **color scatter** curves are the data for five different concentrations of the same interaction. The concentration values are shown in the legend. The **black solid** curves are the fitting results using the bivalent analyte model. The vertical **black dashed** line separates the association and dissociation phases.

In Fig 7, we show violin log-plots for the estimated kinetics parameters for data sets with standard length of dissociation for the bivalent analyte model (see S1 Table for full details). Based on Fig 7, we found that the estimated values for *k*_*a*1_ and *k*_*a*2_ are consistent for most of the data sets after performing global fitting on 5 concentrations for each data set. Most of estimated *k*_*a*1_ values are of order of magnitude 3 and *k*_*a*2_ consistently estimated at about 10^-4^ *RU*^-1^s^-1^ (see S1 Table). On the other hand, the order of magnitude for *k*_*d*1_ values are are less consistent and are estimated to between −3 to −2 (see S1 Table). For *k*_*d*2_, the values for data sets 1 and 3 are estimated to be the lower bound 0 while their standard errors are 3.56 × 10^-6^ *s*^-1^ and 3.11 × 10^-6^ *s*^-1^, respectively (see S1 Table). Furthermore, for each violin log-plot, we provide an associated coefficient of variation (CV), which can be computed by dividing the standard deviation for each parameter by the mean of each parameter across the data sets. The computed CV for *k*_*a*1_, *k*_*a*2_, *k*_*d*1_, and *kd_2_* are 0.20, 0.25, 0.81, and 0.85, respectively. This result suggests that the estimated values for *k*_*a*1_ and *k*_*a*2_ are less disperse compared to *k*_*d*1_ and *k*_*d*2_. This led to our hypothesis that the dissociation rate constants might be practically non-identifiable with the current standard length of dissociation.

**Fig 7:**
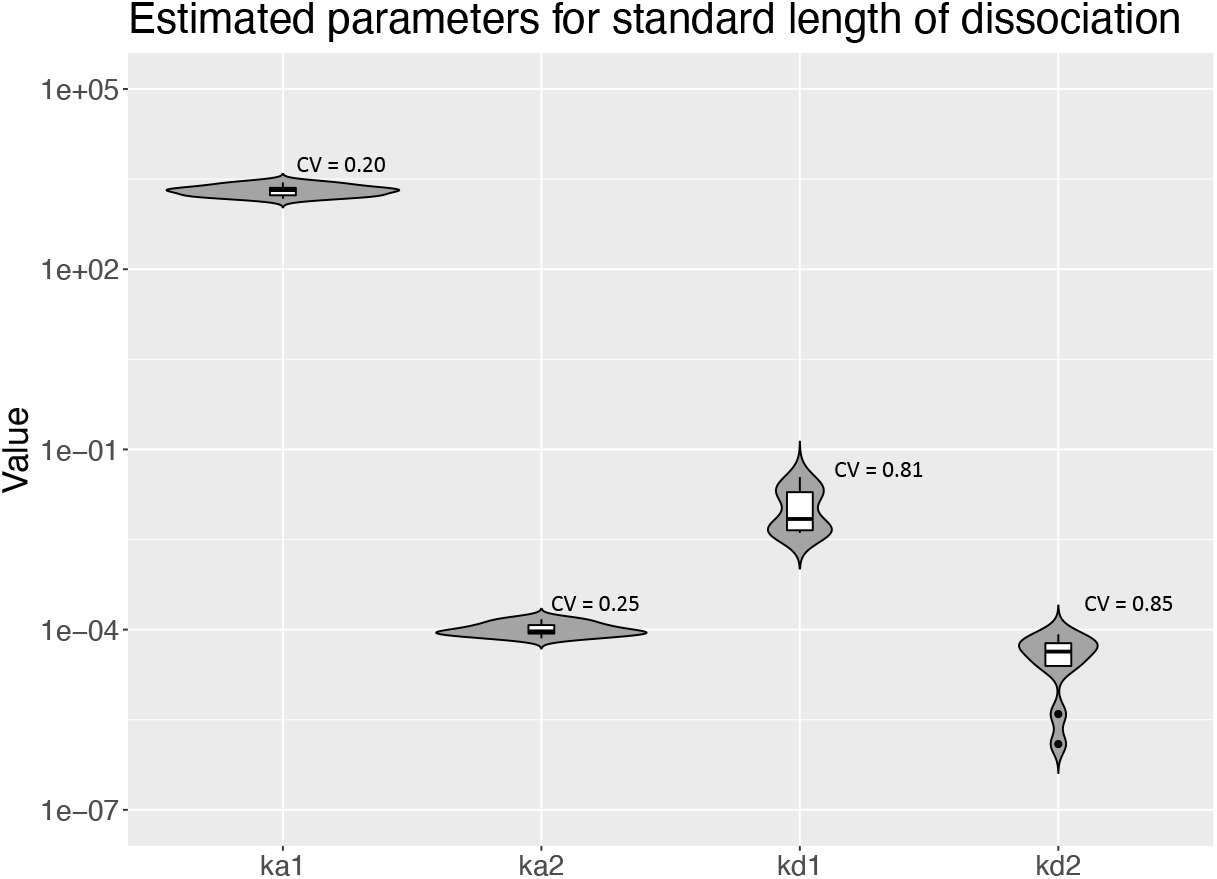
Violin log-plots for estimated parameter of the bivalent analyte model for standard length of dissociation. We illustrate the dispersion of the estimated parameters for the bivalent analyte model for standard length of dissociation using violin log-plots. From left to right, we show the violin log-plots for *k*_*a*1_, *k*_*a*2_, *k*_*d*1_, and *k*_*d*2_. In addition, we provide the computed coefficient of variation (CV) for each parameter. Note that the estimated parameters for data sets 9 and 10 are not included because the recovered dynamics are 1:1 Langmuir interactions. The estimated values for *k*_*d*2_ for datasets 1 and 3 are also excluded since log_10_(0) = −∞.

### 5.2 Log profile likelihood identifies *k*_*d*2_ as a non-identifiable parameter

As mentioned in Section 5.1, we hypothesized that one or both of the dissociation parameters might be practically non-identifiable with the standard length of dissociation. To investigate our hypothesis, we carried out parameter identifiability analysis using the log profile likelihood method as described in Section 4.4. First, we simulated synthetic data for the standard length of dissociation, i.e., 600 seconds. The parameter values used to generate synthetic data are displayed in Table 2.

**Table 2:** Parameter values for simulations. Parameter values used for simulation to study parameter identifiability analysis. Using these values, we generated synthetic noisy data with standard (600 seconds) and extended (1780 seconds) length of dissociation.

| Parameter | $k_{a1}$ | $k_{d1}$ | $k_{a2}$ | $k_{d2}$ | $R_{\max_{1,\dots,5}}$ | $t_{1,\dots,5}^*$ |
| --- | --- | --- | --- | --- | --- | --- |
| Value | $2.00 \times 10^3$ | $4.00 \times 10^{-3}$ | $5.00 \times 10^{-5}$ | $1.00 \times 10^{-5}$ | 600 | (283, 300, 282, 222, 145) |

To simulate experimental data, we added normally distributed noise, 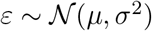 with *μ* = 0 and 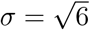. We note that all these values are chosen based on the results of parameter estimation on the experimental data with the standard length dissociation. In addition, *σ*^2^ = 6 corresponds to 2% to 6% of the maximum response depends on the sensorgram. This satisfies the recommendation of having residual values being less than 10% of the maximum response of the fitted curved for a quality fit [24]. After generating synthetic noisy data, we analyzed the identifiability for kinetics parameters, *k*_*a*1_, *k*_*a*2_, *k*_*d*1_ and *k*_*d*2_. In Fig 8a, we displayed the computed profile likelihood for *k*_*d*2_ with standard length of dissociation. In this figure, *k*_*d*2_ profile resembles the practically non-identifiable example being shown in Fig 5a, i.e., the profile is shallow on one side, particularly, on the left side in this case. This means, regardless of *k*_*d*2_ values on the left side, the algorithm is almost always able to achieve the global minimum error. This observation further explains our results for estimated values of *k*_*d*2_ in S1 Table.

**Fig 8:**
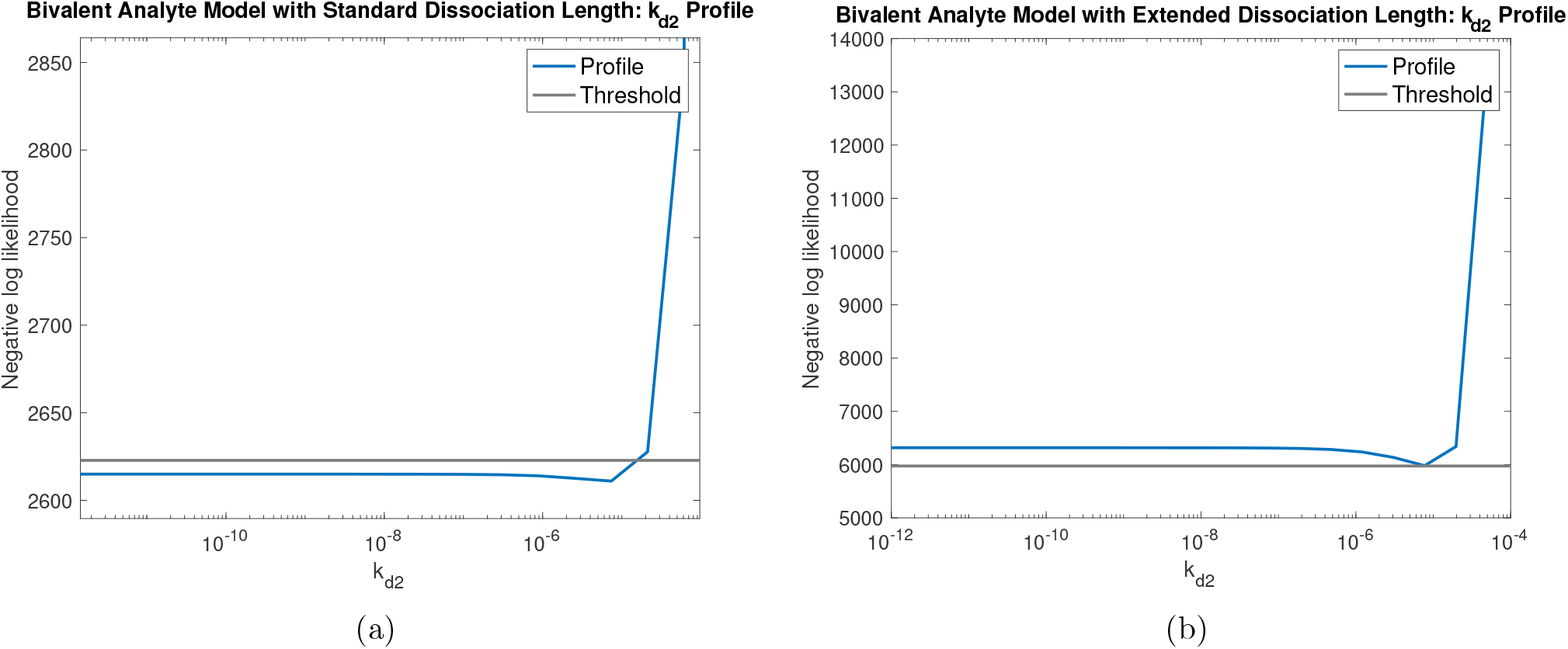
Parameter identifiability analysis on synthetic noisy data. Profile likelihood for *k*_*d*2_ with: (a) **standard** length of dissociation and (b) **extended** length of dissociation. For **standard** length of dissociation (a), the profile for *k*_*d*2_ resembles an L-shape with the left side stays below the threshold. This indicates that *k*_*d*2_ is not practically identifiable with the standard length of dissociation. For the **extended** length of dissociation (b), the left side of the profile for *k*_*d*2_ stays above the threshold indicating that *k*_*d*2_ is practically identifiable.

On the other hand, the profiles for the remaining kinetics parameters, *k*_*a*1_, *k*_*a*2_, and *k*_*d*1_, manifest the bowl shape example in Fig 5b (See S2 Fig). These profiles reinforce the observation that the estimated values for *k*_*a*1_, and *k*_*a*2_ are stable. While *k*_*d*1_ is somewhat unstable, it does meet the threshold for practical identifiabilty, and the highly variable *k*_*d*2_ is not identifiable.

To further reinforce our results, we simulated two sets of noisy synthetic data with the standard length of dissociation with *k*_*d*2_ = 10^-5^ *s*^-1^ and *k*_*d*2_ = 0 *s*^-1^. Note that all other parameters were kept the same as described in Table 2. In Fig 9a, the fuzzy red curves represent the synthetic noisy data with *k*_*d*2_ = 10^-5^ *s*^-1^ for 5 concentrations while the scatter blue curves correspond to the synthetic noisy data with *k*_*d*2_ = 0 *s*^-1^. Even though the two values for *k*_*d*2_ are on different orders of magnitude, the dissociation phases are almost identical in the presence of noise in Fig 9a. Based on these results, we concluded that *k*_*d*2_ is practically non-identifiable with the standard length of dissociation.

**Fig 9:**
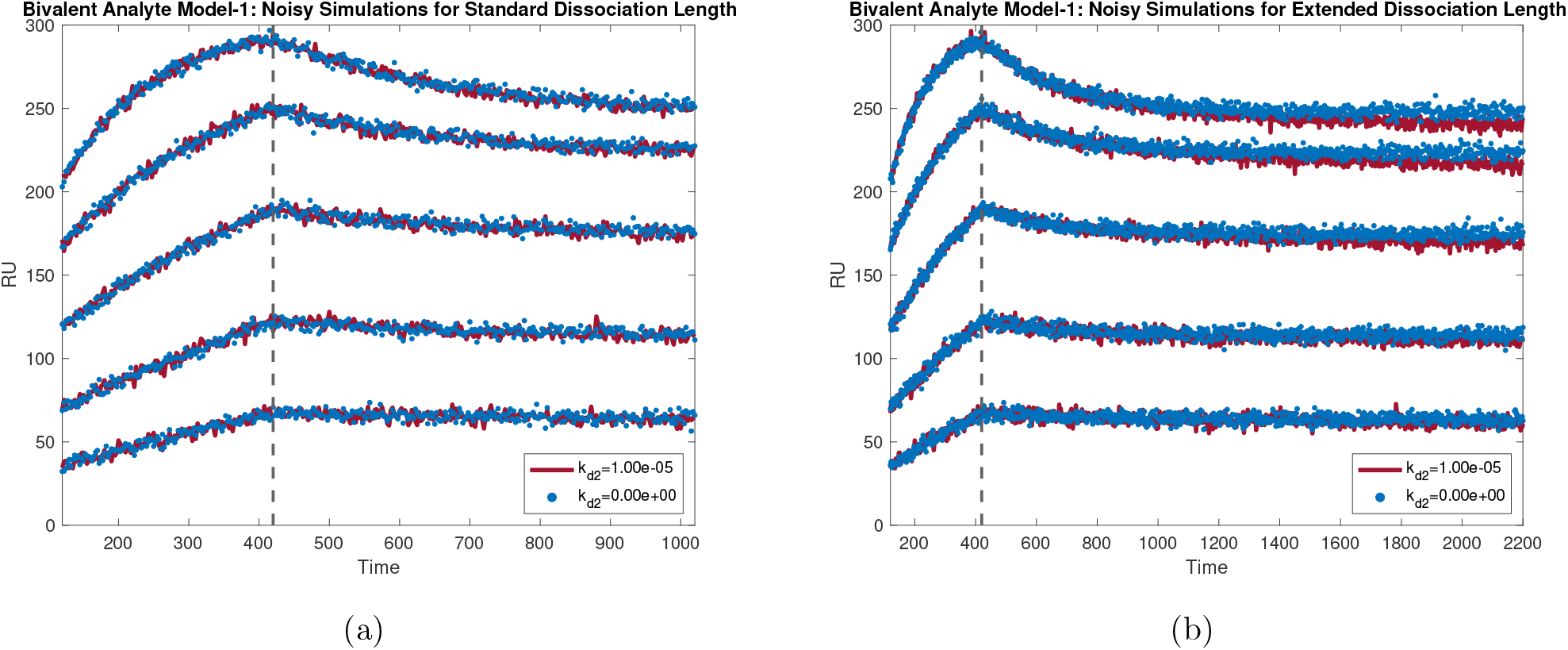
Comparison of noisy simulated synthetic data. Comparison plots of noisy simulations for: (a) **standard** dissociation length and (b) **extended** dissociation length. Each figure shows the noisy simulated solutions of the bivalent analyte binding model with *k*_*a*1_ = 2.00 × 10^3^ *M*^-1^ *s*^-1^, *k*_*d*1_ = 4.00 × 10^-3^ *s*^-1^, *k*_*a*2_ = 5.00 × 10^-5^ *RU*^-1^ *s*^-1^, and two different values for *k*_*d*2_: (**red**) 10^-5^ *s*^-1^ and (**blue**)0 *s*^-1^. For the **standard** length of dissociation (a), the noisy simulated solutions for both *k*_*d*2_ = 10^-5^ *s*^-1^ and *k*_*d*2_ = 0 *s*^-1^ are indistinguishable. On the other hand, for the **extended** dissociation length (b), the noisy simulated solution of the bivalent analyte binding model with *k*_*d*2_ = 10^-5^ *s*^-1^ can be distinguished from the same model solution with *k*_*d*2_ = 0 *s*^-1^.

### 5.3 Simulation provides optimal experimental conditions

A source of practical non-identifiability is lack of sufficient information in the data. As suggested in previously published literature [18, 20, 29], one approach is to collect additional data so that there is sufficient information for the algorithm to uniquely estimate parameters. While it is not possible to collect more data by increasing the sampling frequency due to the limitation of our equipment, collecting additional data by increasing the length of dissociation phase is achievable. Therefore, we examined parameter identifiability on the same synthetic data, but with extended length of dissociation phase. Here, we chose the length of dissociation to be 1780 seconds. The profile likelihood for *k*_*d*2_ is shown in Fig 8b. Identifiability analysis showed that with the extended length of dissociation, *k*_*d*2_ is practically identifiable as both sides of *k*_*d*2_’s profile are non-flat according its corresponding computed threshold. We again further reinforced our conclusion by simulating synthetic noisy data again for two different *k*_*d*2_ values, *k*_*d*2_ = 10^-5^ *s*^-1^ (fuzzy red curves) and *k*_*d*2_ = 0 *s*^-1^ (scatter blue curves), with extended length of dissociation. In Fig 9b, we displayed the comparison for such simulations. We found that the distinction between the two simulated noisy synthetic data are much more noticeable compared to the case with standard length of dissociation. For other kinetics parameters, their profiles preserve their practical identifiability as shown in S3 Fig 9.

In conclusion, all kinetics parameters, including *k*_*d*2_, are practically identifiable with extended length of dissociation.

### 5.4 Kinetics parameters can be reliably estimated with extended length of dissociation

As demonstrated in Section 5.2, all kinetic parameters for the bivalent analyte model are practically identifiable with the extended length of dissociation using noisy synthetic data. In this section, we discussed our parameter estimation results on experimental data with extended length of dissociation. In Fig 10, we show the results for the first data set as a representative example (see S2 Fig for all 14 replicates). Similar to the results in Fig 6 (see S1 Fig for all 14 replicates) with the standard length of dissociation, the bivalent analyte model is able to describe the interaction between CH31 mAb and CH505 T/F gp120 (see S2 Fig for all 14 replicates).

**Fig 10:**
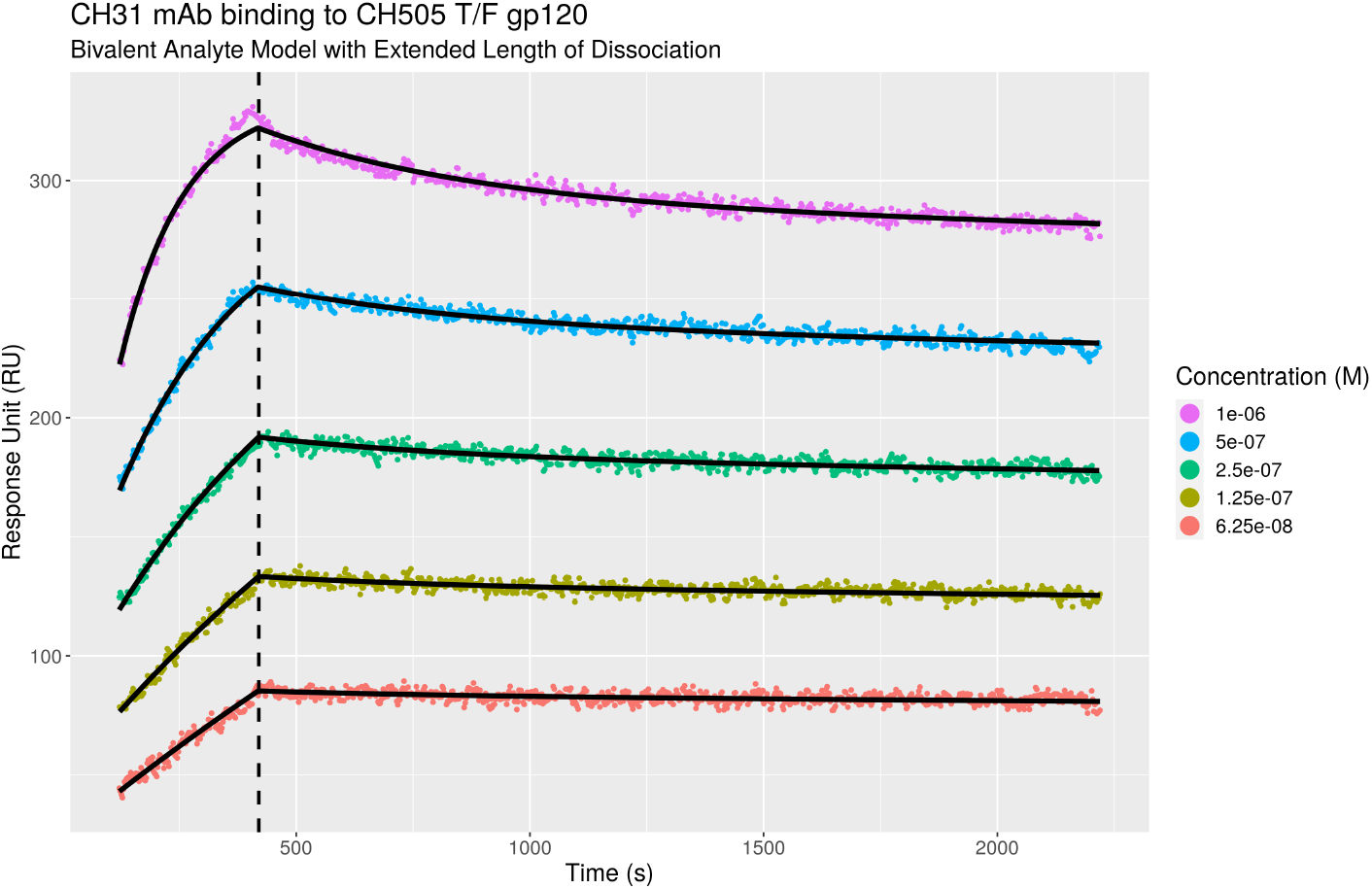
A representative bivalent analyte model fitting result with extended length of dissociation. Bivalent analyte model fitted sensorgrams of CH31 mAb binding to CH505 T/F gp120 are shown. The **color scatter** curves are the data for five different concentrations of the same interaction with the concentration values are shown in the legend. The **black solid** curves are the fitting results using the model. The vertical **black dashed** line separates the association and dissociation phases.

As shown in Fig 11, the values for *k*_*d*2_ of the bivalent analyte model with the extended length of dissociation are more consistently estimated compared to the results with the standard length of dissociation Fig 7 (also, see S1 Table and S2 Table). In addition to the improvement in *k*_*d*2_ identifiability, the extended dissociation length improves the consistency of estimates of *k*_*d*1_ as shown in Fig 11. Furthermore, the computed CV for *k*_*d*1_ and *k*_*d*2_ decrease from 0.81 to 0.31 and 0.85 to 0.61, respectively.

**Fig 11:**
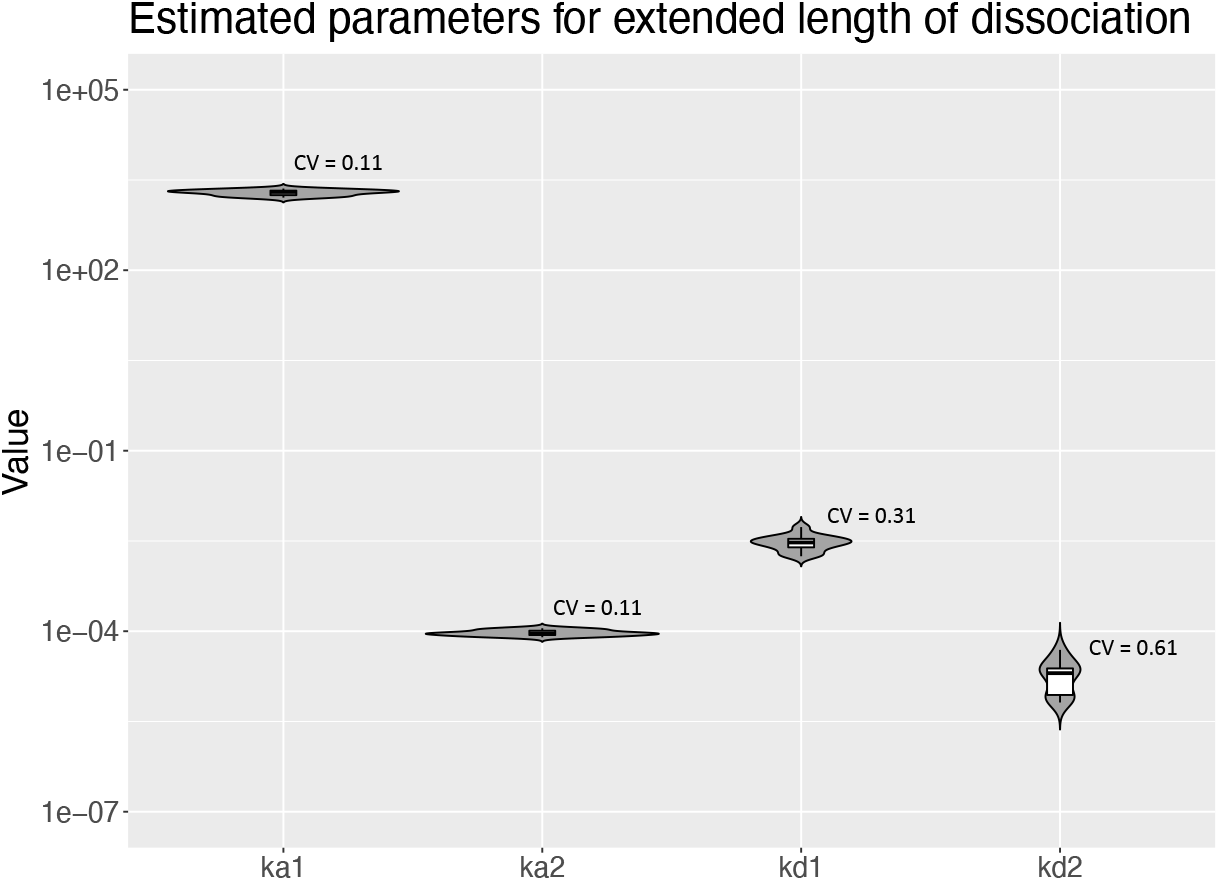
Violin log-plots for estimated parameter of the bivalent analyte model for extended length of dissociation. We illustrate the dispersion of the estimated parameters for the bivalent analyte model for extended length of dissociation using violin log-plots. From left to right, we show the violin log-plots for *k*_*a*1_, *k*_*a*2_, *k*_*d*1_, and k_d2_. In addition, we provide the computed coefficient of variation (CV) for each parameter. Note that the estimated parameters for all data sets are included.

Furthermore, by extending the dissociation length, *k*_*d*2_ can be reliably estimated to be about 10^-5^ as shown in S2 Table. We also see the improvement in consistency in estimation of other kinetics parameters. For example, in S1 Table, *k*_*d*1_ values are estimated to be between 10^-3^ *s*^-1^ to above 10^-2^ *s*^-1^ for data sets with the standard length of dissociation. In contrast, for the extended length of dissociation, *k*_*d*1_ are consistently estimated to be approximately 3 × 10^-3^ *s*^-1^ (see S2 Table).

### 5.5 Importance of addressing the local minima problem

Because the `LM` algorithm is a local optimization algorithm, it is prone to getting stuck in a local minimum when solving non-linear problems and therefore outcomes can depend heavily on the initial starting values of parameters. Therefore, for a robust fit, we performed parameter estimation at multiple sets of initial guesses. For each kinetic parameter (*k*_*a*1_, *k*_*d*1_, *k*_*a*2_, and *k*_*d*2_), we implemented a grid search approach, where we examined all possible combinations of 3 different initial values for each of the four kinetics parameters, while using the same initial values for non-kinetic parameters. This results in 3^4^ or 81 different sets of initial guesses. After running parameter estimation at different initial guesses, we recorded the optimized parameters with the lowest error. Although this approach is computationally expensive, it ensures that recorded optimized parameters give the lowest possible sum of squared error, reducing the chance that the estimated parameters are not optimal. In Table 3, we showed a representation of the grid search outcome for the initial guesses for kinetics parameters for the first dataset of the bivalent analyte model. With “wrong” initial guesses, the algorithm could fail to achieve the global minimum error. For example, with 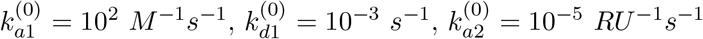, and 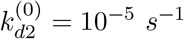 the `LM` algorithm is only able to converge locally. In contrast, if we change the initial guess 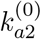 to 10^-4^ *RU*^-1^ *s*^-1^, the algorithm is able to reach the global minimum error. Therefore, when using a local optimization algorithm such as the `LM` algorithm, it is crucial to test the results at multiple initial guesses to ensure a robust fitting result.

**Table 3:**
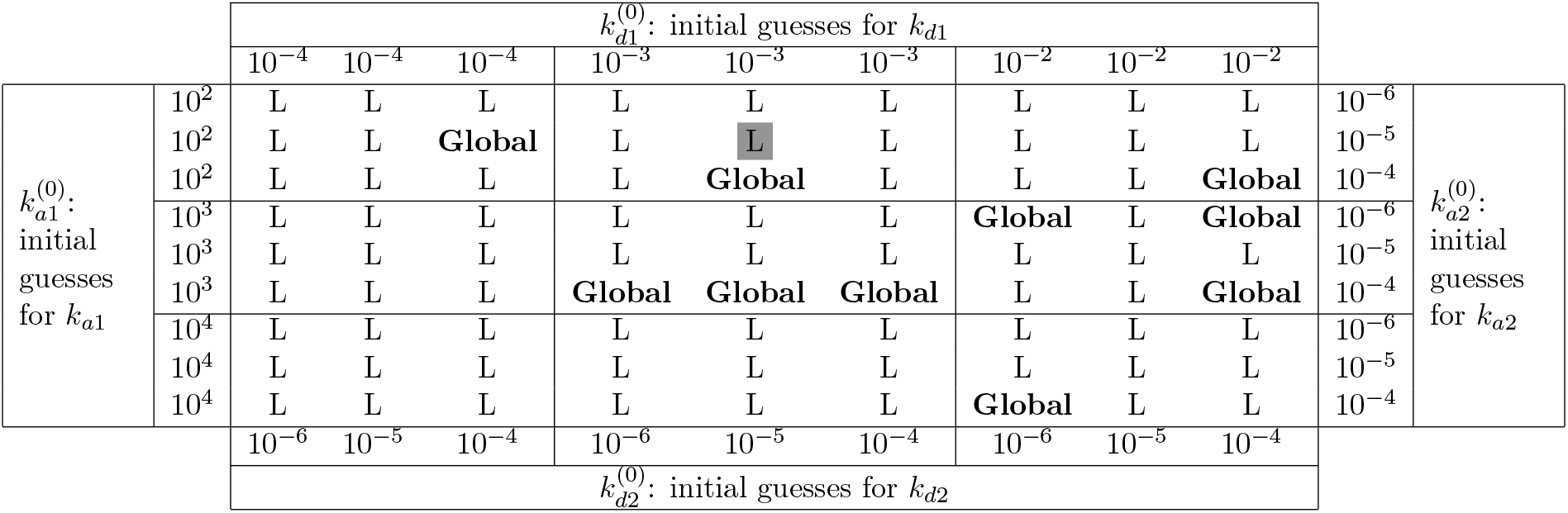
Grid search results table for initial guesses. Table for initial guesses grid search results for a representative dataset for the bivalent analyte model with standard length of dissociation. The outcomes are divided into two categories: L, and **Global**. Given a set of initial guesses, L stands for local minimum, where the `LM` algorithm gets stuck at the local minimum. In contrast, when the `LM` algorithm reaches the global minimum, we denote the result as **Global**. An example to read the table: at the gray-boxed L, with the initial guesses 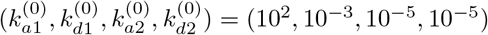, the `LM` algorithm converges to a local minimum.

## 6 Discussion

Biophysical determinations of antibody-antigen interactions directly inform selection of mAbs for immuno-prophylaxis trials and novel immunogen design. For SPR data of antigen binding to immobilized mAbs, the 1:1 Langmuir binding model is appropriate [1, 2, 3, 4]. However, when using mAbs as analyte binding to antigens, a bivalent analyte binding model is required to better describe the kinetics data unless the antigen is immobilized to an optimal density to eliminate avidity effects. Analysis for bivalent analyte models for low-throughput data using commercial software has been done in previous studies [7, 11, 12, 13, 14], but these programs cannot be easily applied to high-throughput, non-regenerative data, and they do not address local minima or parameter identifiability.

In this work, we have introduced a robust parameter estimation pipeline for a bivalent analyte model that can be applied to high-throughput data and that directly addresses the problems of local minima and parameter identifiability. We further used our identifiability analysis to optimize the experimental design. Because the parameter in question was the second dissociation rate constant, we simulated experiments with extended length of the dissociation phase and proposed an optimal duration of observation. We then collected data under the proposed design and succeeded in identifying all of the model parameters. Simulation, together with identifiability analysis can save time, materials and funds, by providing information about experimental variables such as length of dissociation data collection.

We used a common local optimization algorithm, the Levenberg–Marquardt algorithm [28]. Our results from the grid search (see Table 3) show that with the “wrong” (yet physically reasonable) initial guess, it is possible to report results at a local minimum. Different kinetic interactions will have different optimal starting values. We therefore used a grid of 81 initial values. This process is very computationally expensive and inefficient: The full analysis took approximately 4 hours on server running R.1.4.17 [31] using parallel computing with 14 cores and 16 GB RAM. For future work, we will consider methods to better select initial guesses or implement a global optimization algorithm [32, 33].

The parameter estimation and grid search are currently implemented in R [31] and are therefore open source. The profile likelihood analysis is implemented in Matlab. We plan to include the profile likelihood in R to include in our package that is under development.

## 7 Funding

This study was supported by a grant (INV-0008612) for the Antibody Dynamics platform of the Global Health Discovery Collaboratory (GHDC) from the Bill and Melinda Gates Foundation (BMGF) and partly supported by a grant for the Antibody Dynamics platform of the Global Health – Vaccine Accelerator Platforms (GH-VAP) from the BMGF to GDT (OPP12109388), an NIAID education research program (R25AI140495) and the Duke University Center for AIDS Research (CFAR)(5P30 AI064518). KN was supported by the National Science Foundation Graduate Research Fellowship under Grant No. DGE-2137100. Any opinion, findings, and conclusions or recommendations expressed in this material are those of the authors and do not necessarily reflect the views of the funding sponsors.

## 8 Acknowledgements

We would like to thank Dr. Chi Wei Cliburn Chan and the Duke Center for AIDS Research for providing Kyle Nguyen the summer internship opportunity that initiated this work. We acknowledge Dr. Kevin Saunders and Dr. Barton F. Haynes of Duke Human Vaccine Institute for HIV-1 envelope glycoprotein and antibody. We thank Dr. James Peacock in the Duke Human Vaccine Institute Protein Production Facility, which received funding support from the Collaboration for AIDS Vaccine Research from the BMGF (OPP1066832) for protein and antibody production work.

Fig 4 and Fig 3a were created with BioRender.

## 9 Supplemental materials

**S1 Fig. Bivalent analyte model fitting results for the standard length of dissociation.** Model fitting results using the bivalent analyte model for: **(a)-(n)** data sets 1-14 with **standard** length of dissociation.

**S2 Fig. Bivalent analyte model fitting results for the extended length of dissociation.** Model fitting results using the bivalent analyte model for: **(a)-(n)** data sets 1-14 with **extended** length of dissociation.

**S3 Fig. Profile likelihood for kinetics parameters** *k*_*a*1_, *k*_*d*1_, *k*_*a*2_, **and** *k*_*d*2_ **on synthetic noisy data with standard length of dissociation.** Parameter identifiability analysis results on synthetic noisy data with standard length of dissociation. We use negative log profile likelihood as the method to perform parameter identifiability analysis on 4 kinetics parameters: (a) *k*_*a*1_, (b) *k*_*d*1_, (c) *k*_*a*2_, and (d) *k*_*d*2_.

**S4 Fig. Profile likelihood for kinetics parameters** *k*_*a*1_, *k*_*d*1_, *k*_*a*2_, **and** *k*_*d*2_ **on synthetic noisy data with extended length of dissociation.** Parameter identifiability analysis results on synthetic noisy data with extended length of dissociation. We use negative log profile likelihood as the method to perform parameter identifiability analysis on 4 kinetics parameters: (a) *k*_*a*1_, (b) *k*_*d*1_, (c) *k*_*a*2_, and (d) *k*_*d*2_.

**S1 Table. Estimated kinetics parameters for bivalent analyte model for data sets with standard length of dissociation.** Estimated kinetics parameter for the bivalent analyte model with standard length of dissociation.

**S2 Table. Estimated kinetics parameters for bivalent analyte model for data sets with extended length of dissociation.** Estimated kinetics parameter for the bivalent analyte model with extended length of dissociation.

**S1 Appendix. Pseudocode for parameter identifiability.** The appendix contains the pseudocode for parameter identifiability.

**S2 Appendix. Sensorgrams for bivalent analyte with standard length of dissociation.** The appendix contains the sensorgrams for all data sets with standard length of dissociation.

**S3 Appendix. Sensorgrams for bivalent analyte with extended length of dissociation.** The appendix contains the sensorgrams for all data sets with extended length of dissociation.

## Notes

### Competing Interest Statement

The authors have declared no competing interest.

